# Glutamatergic synaptic resilience to overexpressed human alpha-synuclein

**DOI:** 10.1101/2025.02.06.636960

**Authors:** Patrícia I. Santos, Inés Hojas García-Plaza, Ali Shaib, Jeong Seop Rhee, Abed Alrahman Chouaib, Silvio O. Rizzoli, James Daniel, Tiago F. Outeiro

**Affiliations:** Department of Experimental Neurodegeneration, Center for Biostructural Imaging of Neurodegeneration, University Medical Center Göttingen, Göttingen, Germany; Department of Molecular Neurobiology, Max Planck Institute for Multidisciplinary Sciences, Göttingen, Germany; Department of Neuro- and Sensory Physhiology, University Medical Center Göttingen, Humboldtallee 23, 37073, Göttingen, Germany; Department of Cellular Neurophysiology, Center for Integrative Physiology and Molecular Medicine (CIPMM), Saarland University, Homburg, Germany; Center for Biostructural Imaging of Neurodegeneration (BIN), Göttingen, Germany; Translational and Clinical Research Institute, Faculty of Medical Sciences, Newcastle University, Newcastle upon Tyne, United Kingdom; Max Planck Institute for Multidisciplinary Sciences, Göttingen, Germany; Scientific employee with an honorary contract at Deutsches Zentrum für Neurodegenerative Erkrankungen (DZNE), Göttingen, Germany

**Keywords:** alpha-synuclein, Parkinson’s disease, synaptic transmission, autapses, glutamatergic neurons

## Abstract

Alpha synuclein (aSyn) is an abundant protein that, in the brain, is concentrated in neuronal presynaptic boutons and associates with synaptic vesicles. aSyn is strongly linked with Parkinson’s disease (PD) due to its accumulation in pathognomonic inclusions in neuronal cells, including in glutamatergic neurons. While increased expression of wild-type (WT) aSyn due to multiplications of the SNCA gene, or the expression of mutant forms of aSyn, can cause familial forms of PD, it is still unclear whether increased levels of aSyn alone are sufficient to impair synaptic function. Previous studies have used experimental systems that do not allow systematic characterisation of presynaptic physiology. To address this gap in research, we used lentiviral vectors to overexpress human aSyn (haSyn) in continental and autaptic glutamatergic neurons from rodent hippocampus and systematically analysed their presynaptic function. Virally-transduced neurons exhibited levels of expression of haSyn that mimic those associated with triplications of the SNCA gene in PD patients (2-fold increase in total aSyn), as determined using quantitative immunofluorescence imaging and immunoblots. Neuronal toxicity, neuronal morphology, and SNAP-25, a presynaptic protein, were not altered in continental cultures. Finally, a systematic characterization of autaptic neurons expressing haSyn exhibited no significant difference in any parameter of synaptic function, including basal properties of evoked and spontaneous neurotransmitter release, and synaptic plasticity compared to neurons infected with a control virus. These results indicate that rodent glutamatergic neurons are resilient to aSyn overexpression. In conclusion, our findings suggest neurotoxicity associated with aSyn overexpression is not universal, and that a deeper understanding of aSyn biology and pathobiology is necessary.

## INTRODUCTION

Alpha- synuclein (aSyn) is a soluble 14.5 kDa protein expressed in the neurons of vertebrates ^1^, and is concentrated in presynaptic boutons, and in the nucleus ^1–3^. Given the concentration of aSyn at presynaptic boutons and its ability to interact directly with the membrane of synaptic vesicles (SVs), it has been presumed that aSyn regulates presynaptic function ^4–6^.

aSyn is a key player in context of the neurodegenerative illness such as Parkinson’s disease (PD) and dementia with Lewy bodies (DLB) due to its accumulation in intracellular protein aggregates known as Lewy bodies and Lewy neurites, composed primarily of aSyn ^7,8^. Another typical neuropathological hallmark of PD is the loss of dopaminergic neurons in the substantia nigra pars compacta, causing dopamine (DA) depletion in the dorsal striatum and various characteristic neurological symptoms ^9^. While the association between aSyn and PD is well established, the precise mechanisms by which the protein may impact neuronal function and cause neurotoxicity remain somewhat enigmatic in spite of extensive research ^4–6,10^. Genetic ablation of aSyn, either alone or in combination with other synuclein family members (β and γ), results in very mild and variable results on synaptic transmission within the glutamatergic system of the brain ^11–16^. Thus, while aSyn is abundant at presynaptic boutons it does not appear to be essential for synaptic transmission or plasticity, at least in experimental models.

Certain familial forms of PD are associated with duplications or triplications of SNCA, the gene that encodes for aSyn gene, resulting in increased aSyn protein expression ^17^. This has led to a proliferation of cellular and rodent disease models in which human aSyn (haSyn) is overexpressed, with the intention of gaining insight into both the physiological function of aSyn and its role in disease. However, the impact of over-expressing haSyn on synaptic function has varied considerably between studies. In glutamatergic neurons, the overexpression of haSyn did not alter neuronal morphology, but was reported to cause various functional changes such as impaired basal glutamatergic neurotransmission ^18^, altered spontaneous neurotransmission ^14,18,19^, altered short-term plasticity ^13,20^, altered long-term potentiation ^13^, slowed kinetics of SV exocytosis ^18,21^, and abnormal SV dynamics ^18,22^. These inconsistencies are likely due to the different model systems and experimental approaches used. Importantly, none of these studies employed an experimental system in which presynaptic function in glutamatergic neurons can be systematically dissected in terms of the pre- and postsynaptic components of neurotransmission. An ideal system for interrogating synaptic function is the autaptic neuronal culture, in which single neurons are cultured in isolation. Within this system, a range of fundamental parameters of synaptic function can be examined independently, allowing a diagnostic approach to studying the impact of molecular changes on the level of a single neuron. Using mild lentivirus-mediated overexpression of haSyn to mimic the levels associated with triplications of the SNCA gene in humans, we examined a range of functional parameters of synaptic function in autaptic glutamatergic neurons with increased aSyn expression. haSyn trafficked to synapses in rodent neurons and caused an overall increase in the total level of aSyn in presynaptic boutons. This aSyn overexpression did not induce cytotoxicity or alter neuronal morphology. Importantly, we observed no effect of aSyn overexpression on SV fusion, synaptic plasticity or endocytosis as studied using whole cell patch clamp electrophysiology. We conclude that ‘mild’ haSyn overexpression does not induce detectable synaptic dysfunction in an autaptic model system. These data indicate that glutamatergic neurons are more resilient than dopaminergic neurons to the expression of increased levels of haSyn.

## MATERIALS AND METHODS

### Primary Hippocampal Neuronal Cultures from Rats

Preparation of primary hippocampal neuronal cultures from E18 Wistar rat embryos were carried out as previously described with slight modifications ^23,24^. All animal procedures were performed in accordance with the European Community (Directive 2010/63/EU), and in compliance with protocols approved by institutional and national ethical committees (Landesamtes für Verbraucherschutz und Lebensmittelsicherheit (LAVES), Braunschweig, Lower Saxony, Germany, license number: T20.7 and 19.3213). In detail, pregnant rats were sacrificed by CO2 inhalation and the embryos were extracted from the uterus irrespective of their sex. The hippocampi were dissected and transferred to ice-cold 1x Hanks balanced salt solution (CaCl2 and MgCl2 free; HBSS) (Gibco) supplemented with 0.5% sodium bicarbonate solution (Sigma-Aldrich). After enzymatic digestion with 0.25% trypsin (Gibco) at 37°C for 15 min, the tissue was centrifugated for 5 minutes at 300 g. Next, 1 mL of FBS was added to the tissue, prior to gentle dissociation and centrifugation. The cells were resuspended in pre-warmed neurobasal medium (Gibco) supplemented with 1% penicillin-streptomycin (Pan Biotech), 0.5% GlutaMax and 2% B27 (Gibco). Primary cells were seeded in 24-well microplates (100,000 cells/well) for immunocytochemistry (ICC) and 12-well microplates (400,000 cells/well) for immunoblotting, all coated with poly-L-ornithine (0.1 mg/mL in borate buffer) (PLO; Sigma-Aldrich). Cells were kept at 37°C with 5% CO2 for 14 to 21 days before they were used for experiments, and one-third of the medium was changed every 3-4 days. Cells were infected with lentivirus coding eGFP or haSyn on day 3 (Multiplicity of infection 1). To limit the growth of glial cells, 4 µM cytosine arabinoside (Sigma-Aldrich) was added once to the cultures.

### Primary Autaptic and Continental Hippocampal Neuronal Cultures from Mice

For the preparation of autaptic neuron cultures, hippocampal neurons were derived from postnatal day 0 (P0) wild-type (WT) from C57BL/6N mice, irrespective of their sex. Animals were kept in groups according to European Union Directive 63/2010/EU and ETS 123. Animal health was controlled daily by caretakers and by a veterinarian. Neuronal cultures were maintained in vitro on glial micro-islands and used for electrophysiological characterization and immunofluorescence assays between 10 and 15 days in vitro. Micro-island cultures were prepared as described previously ^25,26^. Astrocytes used for autaptic cultures glial feeder layers were obtained from cortices dissected from P0 WT mice and enzymatically digested for 15 min at 37°C with 0.25% (w/v) trypsin-EDTA solution (Gibco). Astrocytes were cultured and grown for 7-10 days in vitro (DIV) in T75 flasks in DMEM (Gibco) containing 10% FBS and penicillin (100 U/ml)/streptomycin (100 µg/ml). Next, astrocytes were trypsinized and plated at a density of approximately 30,000 cells per coverslip onto 32 mm-diameter glass coverslips that were first coated with agarose (Sigma-Aldrich). Coverslips were next stamped using a custom-made stamp with a solution containing poly-D-lysine (Sigma-Aldrich), acetic acid, and collagen (BD Biosciences) to generate 400 x 400 µm2 substrate islands permitting astrocyte attachment and growth.

Hippocampi from WT P0 mice were isolated and digested for 60 min at 37°C in DMEM containing 2.5 U/ml papain (Worthington Biomedical Corp.), 0.2 mg/ml cysteine (Sigma), 1 mM CaCl2, and 0.5 mM EDTA. After washing, dissociated neurons were resuspended in pre-warmed (37°C) serum-free Neurobasal medium (Gibco) supplemented with B27 (Gibco), Glutamax (Gibco), and penicillin/streptomycin (100 U/ml, 100 ug/ml) and seeded onto micro-island plates containing the same medium at a density of approximately 4,000 neurons per 32 mm coverslip. On day 1 following preparation, neurons were washed twice in warm medium to remove debris and infected with lentivirus encoding eGFP or haSyn. Neurons were allowed to mature for 10–15 days before they were used for experiments, and only islands containing single neurons were examined for electrophysiological recordings.

For the preparation of mass cultures, hippocampal neurons were prepared as described above, except that dissociated hippocampal neurons were seeded in the same serum-free Neurobasal medium onto 60 mm-diameter Petra dishes coated with poly-L-Lysine (PLL) and collagen. Two hippocampi were used per 60 mm-diameter dish. For mass cultures, the culture medium was replaced completely one day after plating, after three washes in pre-warmed neuronal media.

### Lentivirus Production

Two plasmids were used to generate lentiviruses for transducing neurons: pWPI-CAAGS-IRES-eGFP, which mediates the expression of eGFP alone, and pWPI-CAAGS-haSyn-IRES-eGFP, which mediates expression of haSyn under the control of the hybrid CAGGS promoter (a combination of the CMV enhancer, promoter, first exon and intron of the chicken beta-actin gene, with the splice acceptor of the rabbit beta-globin gene). These constructs were previously described ^27^.

HEK293FT cells were transfected using Lipofectamine 2000 (Thermo Fischer Scientific) or calcium phosphate (CaPO4) precipitation method following the manufacturer’s instructions. Lentiviral expression constructs were co-transfected with vesicular stomatitis virus glycoprotein (VSV-G) packing and pCMVdeltaR8.2 or pCMVdeltaR8.9 ^28–30^. 6 h after transfection using Lipofectamine 2000, the medium was removed and replaced with DMEM with 2% FBS, penicillin/streptomycin (100 U/ml, 100 μg/ml), and 10 mM sodium butyrate. 16h after transfection with CaPO4 precipitation, the medium was changed to Panserin (PAN Biotech) supplemented with 1% penicillin/streptomycin (PAN Biotech) and 1% MEM. Cell medium was harvested 40 h later and viral particles concentrated using Amicon centrifugal filters (100 kDa cut-off; Millipore), followed by washing the virus in Neurobasal-A medium twice, and then tris-buffered saline twice. Concentrated lentivirus was snap-frozen and stored at -80°C.

### Electrophysiology

Electrophysiological recordings in cultured neurons were performed at room temperature (RT, 22 °C) as described previously ^25,31^. Autaptic neurons were whole-cell voltage-clamped at −70 mV with an EPSC10 amplifier (HEKA) under the control of Patchmaster 2 (HEKA). Intracellular patch pipette solution consisted of (in mM): 136 KCl, 17.8 HEPES, 1 EGTA, 0.6 MgCl2, 4 NaATP, 0.3 mM Na2GTP, 15 creatine phosphate, and 5 U/mL phosphocreatine kinase (315–320 mOsmol/L, pH 7.4). Extracellular solution contained (in mM) 140 NaCl, 2.4 KCl, 10 HEPES, 10 glucose, 4 CaCl2 and 4 MgCl2 (320 mOsm/L), pH, 7.3. EPSCs were evoked by depolarizing the cell from −70 to 0 mV.

In order to estimate the size of the readily-releasable pool of SVs, two complementary methodologies were used: repeated action potential trains and local application of hypertonic sucrose ^32^. Both methods trigger release of the same pool of SVs, but while the action potential-evoked release is calcium-dependent, the sucrose application is not ^32,33^. For action potential-driven release, a 2.5 s train of action potentials at 40 Hz was delivered to neurons. The cumulative amplitude was then calculated by summing the eEPSC amplitudes for all stimuli. The area under the baseline was then integrated. A line of best fit was calculated by linear regression from the last 1 s of the stimulus train, and the RRP finally estimated by extrapolating the line of best fit to time = 0 s. The slope of the line of best fit was also used as a measure of the SV replenishment rate. For the readily-releasable pool size estimate using hypertonic sucrose application, 0.5 M sucrose (in extracellular solution) was applied to the neuron for 6 s using a fast-flowing micro-pipe. Sucrose triggered a 2-3 s long transient EPSC, followed by a steady-state current (in which RRP refilling and release reach equilibrium). The RRP size was then estimated with the steady-state current set as baseline (i.e., not correcting for SV pool replenishment) and measuring the area under the curve, thereby measuring the total charge transferred by the release of the RRP. Vesicular release probability (Pvr) was calculated by dividing the charge transfer mediated by a single action potential-evoked EPSC by the charge transfer during the sucrose response.

Miniature EPSCs (mEPSCs) were measured by recording spontaneous activity in the neuron for 100 s in the presence of 300 nM tetrodotoxin (TTX, Tocris Bioscience). mEPSCs were detected using a template for mEPSC analysis (Synaptosoft) after filtering the trace at 1 kHz. Short-term plasticity was evaluated by recording eEPSCs during a train of 50 depolarizing stimuli delivered at 10 Hz or 100 stimuli at 40 Hz. For analysing 10 Hz or 40 Hz trains, the amplitude of each eEPSC was normalized by dividing it by the amplitude of the first eEPSC. The paired pulse ratio (PPR) was then taken as the normalized value of the second eEPSC (i.e., the ratio of the amplitude of the second eEPSC to the first). For calculating the steady state depression, the normalized amplitude of the last 10 eEPSCs in each train were averaged. All traces were analyzed using AxoGraph version 1.5.4 (AxoGraph Scientific).

### SDS-PAGE and Western Blot Analyses

For quantifying aSyn in mass neuronal cultures infected with lentivirus, cultures were prepared in 60 mm dishes and infected as described above. On DIV 14, neurons were washed three times with in PBS and then lysed in 200 µl of Laemmli buffer (100 mM Tris, 4% SDS, 0,2% Bromophenol Blue [Pierce], 20% Glycerol, 200 mM DTT, pH 6.8) or in RIPA buffer (50 mM Tris, pH 8.0, 0.15 M NaCl, 0.1% SDS, 1.0% NP-40, 0.5% Na-Deoxycholate, 2mM EDTA, supplemented with protease and phosphatase inhibitors cocktail (complete TM protease inhibitor and PhosSTOP TM phosphatase inhibitor; Roche). Samples were then sonicated, centrifuged at 13000 rpm, and finally denatured for 5 min at 95 °C. Samples were separated by SDS-PAGE on a 11-12% gel. Proteins were then transferred to nitrocellulose membranes. Membranes were blocked in 5% skim milk or BSA in PBS-T (0.1% Tween-20 in PBS) for 1h and then incubated for 2h or overnight at 4°C with primary antibodies. Afterwards, the cells were washed three times with PBS-T, and then incubated with secondary antibodies for 1 to 2h at RT. Membranes were developed for chemiluminescence using Western Blot HRP Substrate Reagent (GE Healthcare Amersham Hyperfilm) and imaged using an Intas Chemiluminiscence System or Fusion FX Vilber Lourmat, Vilber, France. The membranes that were incubated with fluorescent secondary antibodies were imaged by a fluorescence system (Li-Cor Odyssey® CLx imaging system). The intensity of each band was normalised to beta-actin or beta-tubulin, protein loading controls, and quantified using Fiji software (National Institutes of Health).

### ToxiLight Assay

The cytotoxicicity associated with the overexpression of aSyn on neuronal cultures was assessed using the ToxiLight™ bioassay kit non-destructive bioluminescent cytotoxicity assay (Lonza, Rockland), following the manufacturer’s instructions. Briefly, lyophilized adenylate kinase (AK) detection reagent (AKDR) was reconstituted in 20ml assay buffer and left at RT for 15 minutes to ensure complete rehydration. In a luminescence-compatible 96-well plate, 20 µl of culture supernatant was mixed with 100 µl of AKDR in each well, and after 5 minutes, the plate was measured using an Infinite M200 fluorescence plate reader (TECAN).

### Immunocytochemistry

Neurons were fixed at 14 DIV using 4% PFA (in PBS, pH = 7.4) for 10 min at RT. The neurons were then washed 3 times with PBS and then incubated in 50 mM glycine (in PBS) for 10 minutes to quench autofluorescence. Cells were again washed 3 times and then residual fluorescence was further reduced by incubating the samples for 20 min with Image-iT™ FX (4 drops added to the PBS in each well). After washing, neurons were permeabilised/blocked by incubating in 0.2% Triton X-100 and 5% goat serum (in PBS) or 0.1% Triton X-100 (in PBS). Primary antibodies were then diluted in 2.5% goat serum (in PBS) or 3% BSA and applied to samples overnight at 4 °C. After primary antibody application, samples were washed and secondary antibodies were applied for 60 min at RT. Samples were then washed 3 times and mounted on slides using Aqua-Poly/Mount or Mowiol.

### Confocal Microscopy

Confocal microscopy images from neuronal hippocampal cultures were acquired using the Zeiss LSM 800 – Airyscan, Carl Zeiss Microscopy GmbH, with 20x and/or 63x magnification objectives. Samples were excited using 405, 488, and 561 laser lines, pinhole = 1, 2 averaging line-by-line, 0.250 μm thickness Z stacks, and step size of 1 μm (7-10 slices per neuron). For quantification of axonal length 10-12 images were randomly taken out of three independent experiments, and analysed by ImageJ software.

### Image Analysis

To reconstruct connectivity from 3D image stacks acquired by confocal microscopy, we used a semi-automated neuronal tracing plug-in, Simple Neurite Tracer, from FIJI software as previously described ^34^. Briefly, successive points along the dendrites were selected through the z plane and a path between them was created. Branches were created by defining paths that were starting or ending on other paths. Hessian-based analyses were used to improve the speed and accuracy of the tracing ^35^. For dendritic length measurements, a total of ^30-36^ neurons were randomly traced out of three independent experiments, and their dendritic branching pattern was analysed.

For protein quantification, the neuronal images were analysed as described by ^36,37^. In brief, in-house written scripts in IJ1 and Java language for Fiji and ImageJ (NIH) were used to analyze synaptic proteins. The plugin is available from GitHub (https://github.com/AbedChouaib/Synapse-Count). Automatic thresholds were applied to proteins of interests and areas were converted to masks followed by creating region of interest (ROI) selections. Colocalization analysis was carried out where necessary, colocalizing puncta were measured and ROI selections were applied over the raw images to measure signal intensity. Generated data was produced as columns in csv files. An in-house written script extracted the data and organized them into user-friendly spreadsheets.

For synaptic volume measurements, images were processed using a band pass filtering procedure, following the Crocker and Grier algorithm ^38^ using a Matlab routine. The positions of the different objects were determined in each image of a Z stack, and their connectivity within the stack was estimated by determining their presence in consecutive images. Their volumes were then estimated according to their presence in multiple Z slices, and to their apparent size. The volumes are not corrected for possible diffraction effects.

### Statistical Analysis

Statistical analysis was done using Prism (Versions 5, 6 or 7) software (Graph-Pad Software Inc.). All data were checked for normality distribution using the Shapiro-Wilk test. Statistical comparisons between two groups of data were made using two-tailed unpaired Student’s t-test or Mann-Whitney test. Multiple comparisons were determined using one-way analysis of variance (ANOVA) and Kruskal-Wallis test followed by post hoc Tukey’s and Dunn’s multiple comparison tests.

## RESULTS

### Lentivirus-mediated overexpression of haSyn does not alter neuronal viability or morphology

haSyn overexpression is associated with forms of familial Parkinsonism as well as decreased DA release. Analyses of whether haSyn overexpression impacts synaptic function in glutamatergic synapses, which express abundant aSyn, have yielded conflicting results. To assess whether overexpression of haSyn has an impact on synaptic function, a lentivirus mediating the expression of haSyn was used to infect rodent hippocampal neurons. To determine the extent of expression, we performed western blots from continental cultures of rat hippocampal neurons with an antibody that specifically detects total amounts of aSyn, including the endogenous mouse and the exogenous human forms (Figure 1A-C). As expected, transduced neurons exhibited robust immunolabelling for aSyn (Figure 1A and E). On average, haSyn-infected rat neurons showed an increase of 2.2-fold in the total amount of aSyn compared to control (Figure 1B). We made similar observations in terms of the fold-increase total aSyn when continental cultures of mouse hippocampal neurons were infected with these same lentiviruses (Supplementary Figure S1).

**Figure 1.**
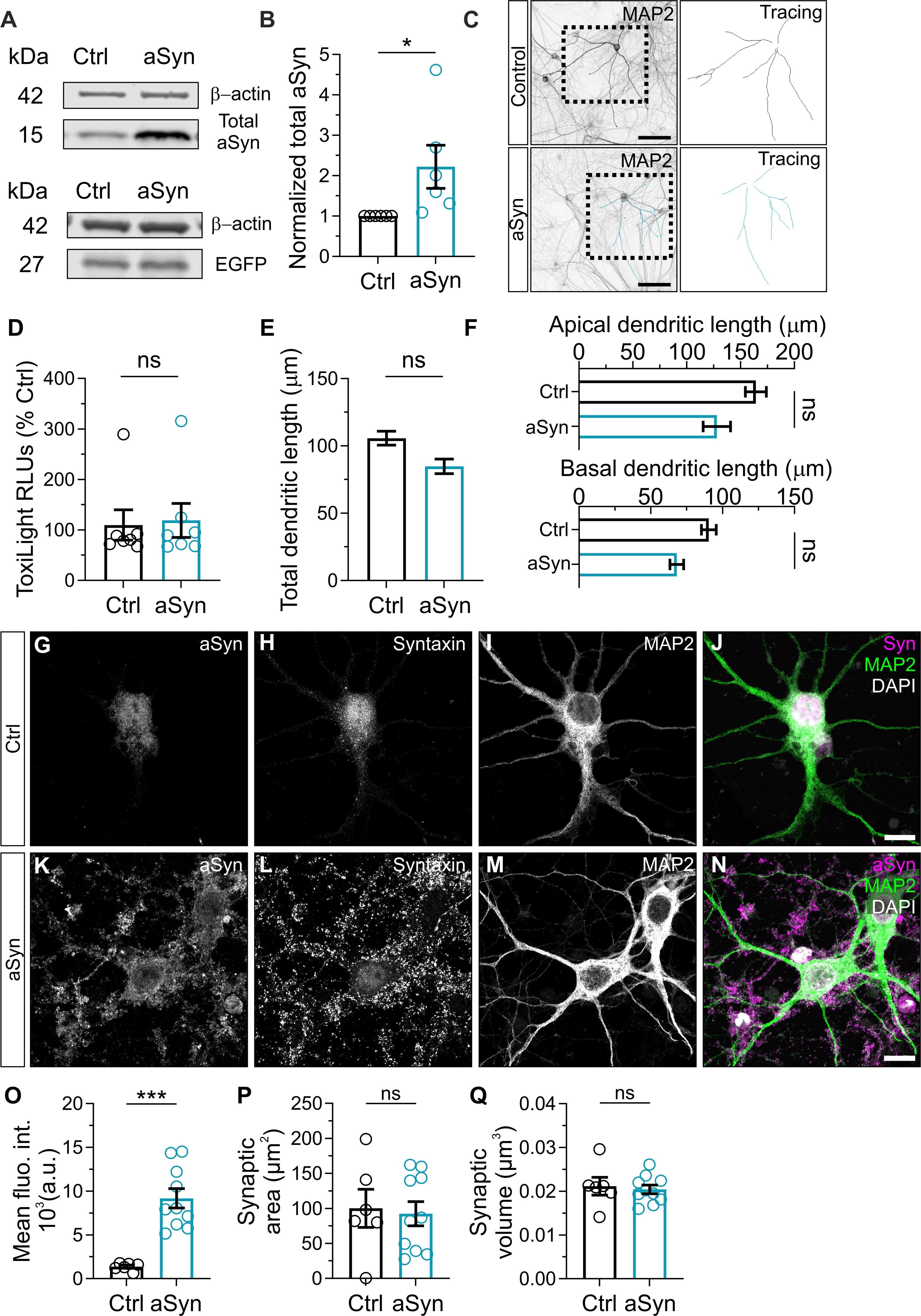
Overexpression of aSyn is induced at hippocampal neurons by infection with haSyn lentivirus. (A-C) Primary hippocampal continental cultures from rat were infected with lentivirus mediating the expression of either eGFP alone or haSyn and eGFP. (A) Representative immunoblots of total aSyn, and eGFP (n=3-6) levels. (B) 2.2-fold increase in total aSyn levels in haSyn-infected neurons (n=6). β-actin was used as a loading control. (C) Representative images of continental hippocampal neurons infected with eGFP or haSyn lentivirus and immunolabelled using anti-MAP2 and corresponding 3D-tracing. (D) Cell death assessed in rat primary neurons transduced with lentiviral encoding for haSyn for up to DIV21 (n=4); haSyn transduction did not alter the total dendritic length (E), apical dendritic length or basal dendritic length (F) at DIV 21 (n=5 neurons from 3 independent experiments per condition. Scale bar = 100 μm. (G-N) Representative images of immunocytochemistry of continental hippocampal neurons infected with haSyn lentivirus and labelled for aSyn (G, K), syntaxin (H,L), and MAP2 (I,M). Overlays are shown (J,N). Scale bar = 10 μm. Quantification of aSyn synaptic mean fluorescence intensity (O), synaptic area (P), and synaptic volume (Q) of neurons infected with haSyn lentivirus compared to neurons infected with eGFP (n=3). All data are expressed as mean ± SEM; Student’s t-test; * p-value ≤ 0.05., *** p-value ≤ 0.001.

haSyn overexpression is associated with neurotoxicity ^39–41^. We therefore investigated whether haSyn overexpression by lentivirus is sufficient to induce neurotoxicity in hippocampal neurons. We used the activity of adenylate kinase in the culture medium as an indicator of membrane integrity and hence neuronal viability. Lentivirus-mediated haSyn overexpression did not induce cytotoxicity compared to eGFP lentivirus-infected neurons (Figure 1D). Thus, these results indicate that increasing total haSyn levels alone does not induce cytotoxicity in cultured hippocampal neurons under the conditions studied.

Several neurological and neurodevelopmental disorders have been correlated with aberrations in dendrite morphology, including alterations in dendrite branching patterns, fragmentation of dendrites, retraction or loss of dendrite branches, anomalous spine density and morphology, and synaptic loss ^42–45^. To assess whether haSyn overexpression has an impact on neuronal morphology, we measured dendritic length in continental cultures of rat hippocampal neurons. Neurons were stained for MAP2 and tracing and measurement of successive branching levels were performed (Figure 1 C). We observed no significant change in total (Figure 1E, apical or basal (Figure 1F) dendritic length in neurons expressing haSyn compared to those expressing eGFP.

To confirm that exogenous haSyn was trafficked to synapses, we used immunocytochemistry. In haSyn-infected continental cultures, we immunolabelled total the presynaptic protein syntaxin to define excitatory presynaptic boutons within neurons (Figure 1H and L) and also immuolabelled total aSyn (Figure 1G, K). The level of aSyn-positive immunolabelling within syntaxin-positive presynaptic boutons was then quantified. A significant increase in mean aSyn grey intensity within presynaptic boutons (Figure 2O) was observed in neurons infected with lentivirus mediating the expression of haSyn, compared to lentivirus mediating the expression of eGFP alone. No differences in presynaptic bouton area (Figure 1P) or volume (Figure 1Q) were observed. An increase in the number of syntaxin-positive puncta was observed in neurons infected with lentivirus mediating the expression of haSyn (Figure 1L).

**Figure 2.**
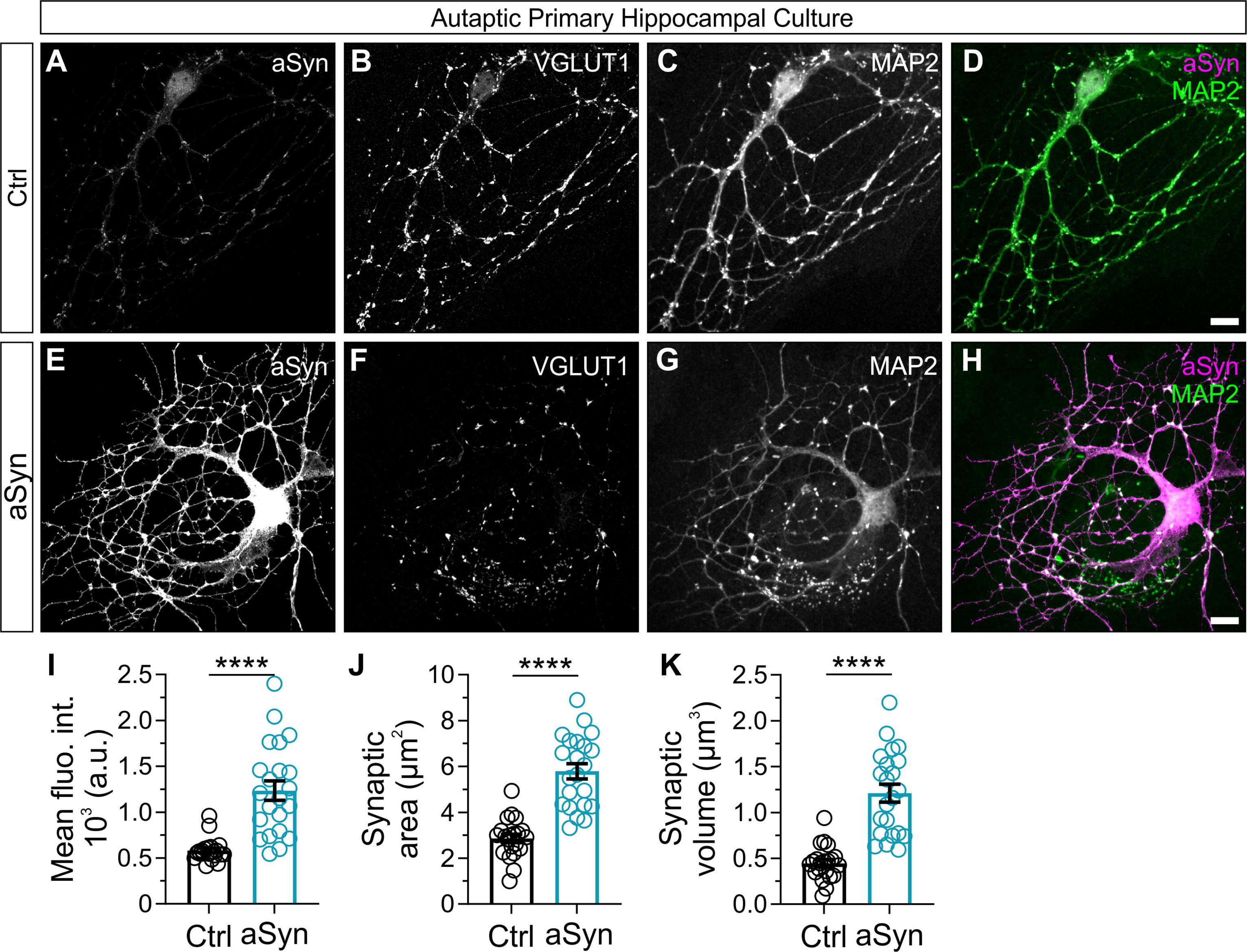
Lentivirus-mediated overexpression of aSyn at synapses of cultured autaptic neurons. (A-H) Representative images of immunocytochemistry of autaptic hippocampal neurons infected with haSyn lentivirus and labelled for total aSyn (A, E), VGLUT1 (B,F), and MAP2 (C,G). Overlays of MAP2 and aSyn are shown (D,H). Scale bar = 10 μm. Column charts showing quantification of aSyn synaptic mean grey intensity (I), synaptic area (J) and synaptic volume (K) for immunolabeled neurons. All data are expressed as mean ± SEM; Student’s t-test; **** p-value ≤ 0.0001.

To examine whether haSyn overexpression has an effect on synaptic architecture or synaptic protein localisation in our continental neuronal cultures, we measured the levels of the presynaptic marker synaptosomal associated protein 25 (SNAP25), as well as the postsynaptic marker postsynaptic density protein 95 (PSD95), through Western blot analysis. Thus, the total expression levels of these synaptic proteins were unaltered by haSyn overexpression (Supplementary Figure S2J-K, and U-V). We also performed immunocytochemistry against SNAP25 (Supplementary Figure S2A-I) and PSD95 (Supplementary Figure S2L-T) to evaluate levels of these proteins at synapses, quantifying the punctate immunofluorescence and estimate the synaptic area and volume. No significant differences were found in the SNAP25 mean grey intensity, area or volume (Supplementary Figure S2D, E and I, respectively) of neurons infected with haSyn when compared with eGFP-infected neurons. In contrast, we observed a significant increase in the PSD95 mean grey intensity and volume (Supplementary Figure S2O and T) between groups. These data indicate that while presynaptic SNAP25 levels were unchanged by haSyn overexpression, the amount of PSD-95 present at postsynaptic structures was elevated when haSyn was overexpressed.

### Overexpression of haSyn does not alter evoked or miniature EPSCs in autaptic glutamatergic neurons

aSyn is a protein predominantly found in presynaptic boutons of neurons, including in glutamatergic neurons ^46^, and it has therefore been presumed to play a role in presynaptic function. To investigate whether the overexpression of haSyn alters glutamatergic neurotransmission we examined synaptic function in autaptic hippocampal neurons infected with lentivirus mediating the expression of eGFP alone or haSyn and eGFP. This model system functions as a highly simplified neuronal circuit and is ideal for dissection and analysis of pre- and postsynaptic properties of neurons.

We first evaluated how infection with lentivirus that mediates expression of haSyn effects the expression level of aSyn at presynaptic boutons compared to infection with lentivirus encoding eGFP alone. In haSyn-infected autaptic cultures, immunolabeling against the glutamatergic SV protein VGLUT1 was used to define presynaptic boutons as subcellular compartments in neurons (Figure 2B and F), and anti-aSyn immunolabelling quantified within these boutons (Figure 2A and E). Across the three metrics examined, mean fluorescence intensity (Figure 2I), synaptic area (Figure 2J) and volume (Figure 2K), synaptic immunolabeling against total aSyn was significantly increased in neurons infected with lentivirus mediating the expression of haSyn, compared to lentivirus mediating the expression of eGFP alone. These data indicate that aSyn was significantly overexpressed at presynaptic boutons from both continental and autaptic cultures infected with haSyn lentivirus.

Synaptic transmission was then analysed in autaptic glutamatergic neurons using whole cell patch clamp electrophysiology, first examining the amplitude of evoked excitatory post-synaptic currents (eEPSCs) as a general measure of synaptic potency. eEPSC amplitude was unchanged in neurons expressing haSyn compared to neurons expressing eGFP alone (Figure 3A and B), indicating that basal synaptic transmission was not affected by haSyn overexpression. To examine whether spontaneous release of glutamate from SVs was influenced by haSyn overexpression, spontaneous miniature EPSCs (mEPSCs) were recorded in infected neurons in the presence of 300 nM TTX. The amplitude and the frequency of mEPSCs were measured (Figure 3C and D, respectively). The amplitude of mEPSCs is a product of both the glutamate content of SVs and the number of post-synaptic glutamate receptors present in the neuron. The frequency of mEPSCs is a product of the probability of SV exocytosis and the number of total synapses formed by the neuron, with a higher frequency indicative of an increased number of synapses. Overexpression of haSyn had no significant impact on the amplitude or frequency of mEPSCs, indicating that the functionality of the postsynaptic receptors, the amount of neurotransmitter, and the number of synapses, were not significantly altered by haSyn overexpression.

**Figure 3.**
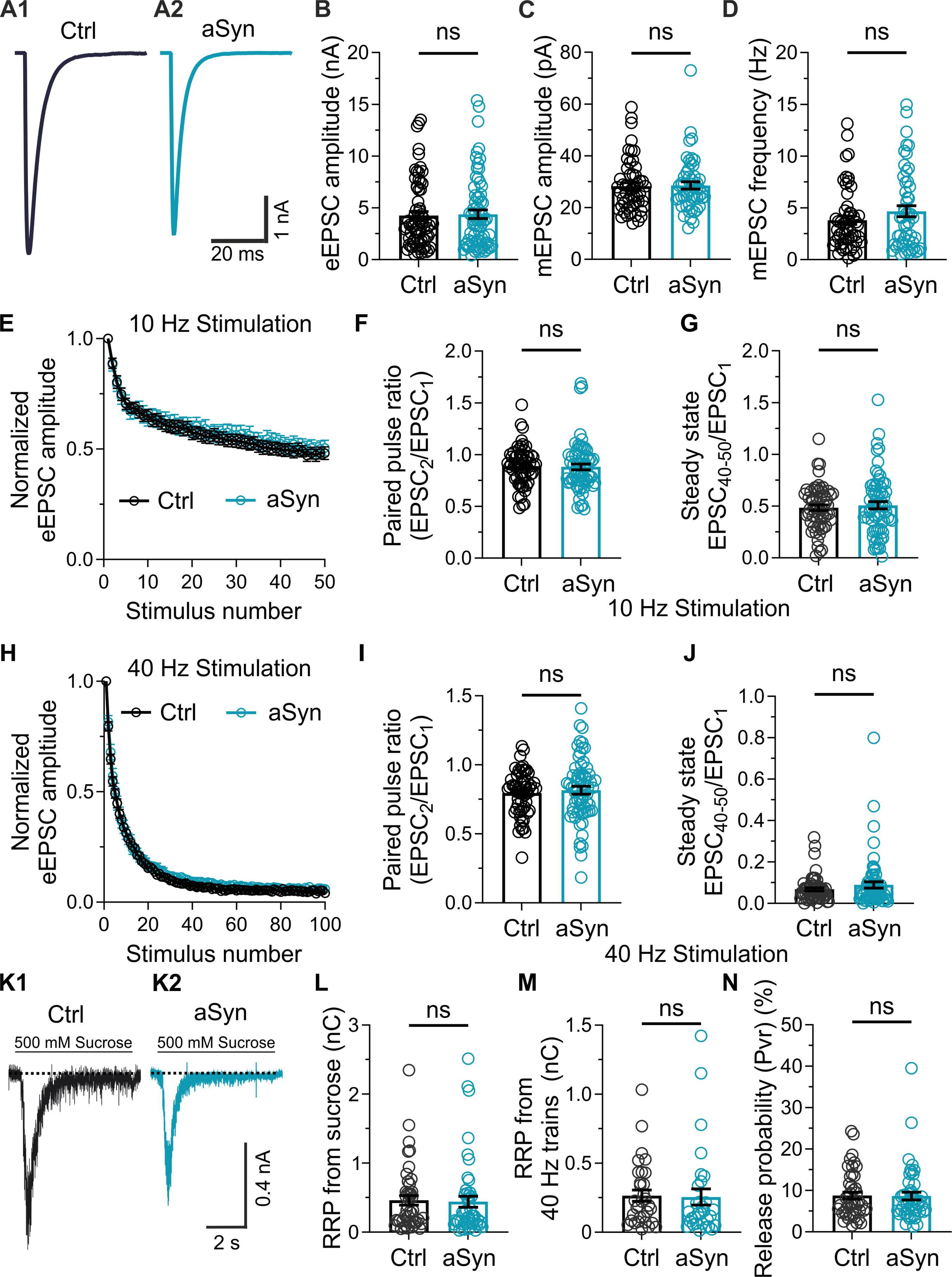
Electrophysiological characterization of functional properties in autaptic glutamatergic neurons over-expressing haSyn. (A) Representative whole-cell patch clamp recordings traces of evoked EPSC amplitudes stimulated at 0.2 Hz from neurons infected with eGFP (A1) or haSyn lentivirus (A2). (B) Quantification of eEPSC amplitudes in neurons expressing eGFP alone (n = 69) or haSyn (n = 70). (C) Quantifications of miniature EPSC amplitudes (C) and frequency (D) recorded in the presence of 300 nM TTX for neurons expressing eGFP (n = 53) or haSyn (n = 52). (E-J) Normalised eEPSC amplitudes from whole-cell patch clamp recordings of neurons stimulated with trains of either 50 APs at 10 Hz (E) or 100 APs at 40 Hz (H) stimulation. Note that the presynaptic plasticity from neurons expressing eGFP or haSyn appeared highly similar, resulting in depression of the eEPSC before reaching a depressed steady state. (F, I) Quantification of the paired pulse ratio (PPR) as the ratio of the first two eEPSCs of each stimulation condition. No differences in PPR were observed between neurons expressing eGFP or haSyn when stimulated at 10 Hz (F) or 40 Hz (I). (G, J) Quantification of the steady-state depression during the stimulation trains, as the average normalised amplitude over the last 10 eEPSCs. For 10 Hz (G), n = 61 for eGFP, n = 67 for haSyn. For 40 Hz (J), n = 59 for eGFP, n = 65 for haSyn. (K-N) Estimation of readily-releasable pool (RRP) size in autaptic glutamatergic neurons over-expressing haSyn. (K) Sample traces showing the current elicited by the application of 500 mM sucrose, representing fusion of SVs in the RRP in neurons expressing eGFP (K1) or haSyn (K2). (L) Quantification of RRP size using the sucrose application method in neurons expressing eGFP or haSyn. (M) Quantification of the RRP estimation based on extrapolating from cumulative eEPSCs evoked by stimulating neurons at 40 Hz. (N) Release probability from neurons expressing eGFP or haSyn. n = 45 for eGFP, n = 48 for haSyn. Data are expressed as mean ± SEM; Student’s t-test or Mann-Whitney test, for normally and not-normally distributed data, respectively.

### Overexpression of haSyn does not alter glutamatergic neurotransmission during high frequency stimulation

Previous studies proposed that haSyn overexpression caused dysfunction in SV dynamics, which could affect SV release during high frequency stimulation. In order to further dissect the mechanisms involved in such a process, we recorded eEPSCs from neurons infected with eGFP or haSyn lentivirus and stimulated with trains of high frequency action potentials. Neurons were stimulated at 10 Hz (Figure 3E-G) and 40 Hz (Figure 3H-J), and the recorded eEPSCs were normalized against the first eEPSC amplitude. At both frequencies, the eEPSC exhibited a stimulation-dependent decrease in amplitude. Normalized eEPSC traces over time appeared similar in neurons over-expressing haSyn compared to eGFP control neurons (Figure 3E and H). Neurons classically exhibit presynaptic plasticity when two depolarizing pulses are applied within a short time frame, such as at the beginning of the trains of repeated stimulation. The second eEPSC typically exhibits either increased (paired pulse facilitation) or decreased (paired pulse depression) amplitude compared to the first eEPSC, depending on the probability of glutamate release at the presynaptic boutons formed by the neuron. The ratio of these two eEPSC amplitudes is referred to as the paired pulse ratio (PPR). Neurons infected with eGFP or haSyn exhibited no significant difference in PPR when stimulated at 10 Hz (100 ms inter-pulse interval, Figure 3F) or 40 Hz (25 ms inter-pulse interval, Figure 3I). During repeated neuronal stimulation, eEPSC amplitudes declined and then eventually reached a steady state. Depression of the eEPSC is mediated by a decline in the number of SVs available for release. Previous research proposed that SV dynamics are impaired when neurons over-express haSyn ^18^, and thus we hypothesized that neurons infected with haSyn lentivirus would exhibit an impaired capacity to maintain neurotransmission during repeated stimulation. To examine this, we calculated the average normalized eEPSC amplitude over the last 10 stimuli of both 10 Hz (Figure 3G) and 40 Hz trains (Figure 3J), which we defined as the depressed steady state. No differences were detected in the depressed steady state between neurons over-expressing haSyn and control neurons at either stimulation frequency. Overall, these data indicate that overexpression of haSyn in glutamatergic neurons did not result in altered presynaptic plasticity or SV dynamics.

### Overexpression of haSyn does not alter the readily-releasable pool of SVs or the vesicular release probability in autaptic glutamatergic neurons

To further characterise presynaptic function in neurons over-expressing haSyn, we examined the size of the readily-releasable SV pool (RRP) in glutamatergic neurons. RRP size is a key factor in synaptic strength ^47^, and is determined both by the overall number of SVs in a presynaptic bouton ^48^ as well as the proteins that facilitate SV docking at the active zone ^49^. We measured the size of the RRP firstly by applying 500 mM hyperosmotic sucrose and measuring the total charge transfer elicited (Figure 3K). No significant difference in RRP size as estimated by sucrose application was observed between neurons over-expressing haSyn and control neurons (Figure 3L). We also calculated the vesicular release probability (Pvr) by dividing the charge resulting from a single eEPSC by the charge transferred during sucrose application. Pvr was no different between neurons over-expressing haSyn and control neurons (Figure 3N). RRP size can also be calculated from the normalized eEPSC plots from neurons stimulated at 40 Hz (Figure 3M). Once again, we found no difference between neurons over-expressing haSyn and control neurons expressing eGFP alone. These data indicated that RRP size and vesicular release probability were unchanged by the overexpression of haSyn.

Overall, the data presented in this study indicate that ‘mild’ overexpression of haSyn has no impact on neuronal viability, morphology, or synaptic function. Importantly, our study constitutes the most comprehensive analysis of synaptic physiology conducted do date, and shows that a significantly increased aSyn levels presynaptic boutons has no impact on pre- or postsynaptic function.

## Discussion

The intraneuronal accumulation of aSyn is a neuropathological hallmark of Lewy body diseases (LBDs) such as PD and DLB, but the mechanisms leading to neuronal dysfunction and death in these disorders are still elusive ^50–52^. Increased levels of aSyn, as observed in familial forms of PD associated with multiplications of the SNCA gene ^17^ are thought to be required for neurotoxicity, although it is unclear whether aSyn overexpression alone is sufficient to induce functional alterations in neuronal activity. Thus, understanding the intricate relationship between aSyn and synaptic function is crucial for developing targeted therapies for LBDs. In this study, we dissected the effects of haSyn over-expression using excitatory hippocampal rodent neurons. Strikingly, we observed no major changes in cytotoxicity, neuronal morphology, or synaptic transmission, including synaptic vesicle exocytosis and endocytosis.

The synaptic function of aSyn remains ambiguous and controversial despite extensive research. Mice lacking aSyn exhibited minimal effects on synaptic transmission, most likely through compensatory mechanisms by other members of the synuclein family ^11,14,53,54^. However, even mice lacking all three forms of synuclein demonstrate only mild alterations in synaptic activity (CITE). Studies involving haSyn overexpression have revealed inconsistent alterations in either SV docking and fusion, endocytosis, and SV dynamics ^18,55–59^. One of the main advantages of the autaptic system is the unprecedented access into presynaptic dynamics, and the potential to elucidate unresolved discrepancies in the field regarding the presynaptic dysfunctions associated with aSyn overexpression. Autaptic neuronal culture allows the analysis of presynaptic and postsynaptic function, short-term plasticity, and synaptic vesicle pool sizes, and this model system has been instrumental in the systematic analysis of the molecular bases of pre- and postsynaptic function in excitatory neurons over the last three decades ^26,37,60,61^.

Based on previous studies, we expected to observe a reduced release of neurotransmitter and defects in the ability to maintain normal synaptic transmission. However, the eEPSC amplitude did not change in human aSyn overexpressing neurons, indicating no differences in basal synaptic transmission. These findings are consistent with other studies investigating overexpression of aSyn in the hippocampus in hippocampal slices ^13,62^, but differ from those in another report that described functional differences in hippocampal primary cultures ^18^. Our results also revealed no difference in Pvr values ^62^, no changes in short-term plasticity (which could have been expected from defects on endocytosis), or in the size of the readily-releasable pool of vesicles ^18^ between neurons over-expressing haSyn and control neurons. Overall, we found that over-expression of h aSyn in glutamatergic hippocampal neurons did not alter the basal amplitude of synaptic evoked fusion events, amplitude or frequency of spontaneous SV fusion events, and presynaptic short-term plasticity. Our results suggest that excess of soluble monomeric haSyn, per se, does not alter its physiological function, and that such alterations may require the accumulation of oligomeric and/or fibrillar forms of haSyn.

The differences between the outcomes of this study with respect to other published data could be due to several factors that varied among different studies. One critical factor is the level of aSyn expression in the experimental model. Here, we over-expressed aSyn up to ∼ 2 -fold. In addition, we showed that, at these levels, aSyn is properly trafficked and present in synapses, and that membrane-integrity is preserved. In contrast, other investigations reported somewhat higher (2.5 to 3-fold) increases in aSyn expression in their models ^18,19^. It is possible that these models result in the accumulation of different aSyn species, which could explain the functional differences observed in these previous studies compared to ours. We also note that we are the first study to use quantitative immunofluorescence to specifically determine the level of aSyn at presynaptic boutons, which is of particular relevance when analyzing synaptic function.

The specific neuronal subtypes and culture conditions used in our study may also impact on the outcomes of aSyn overexpression, as different neuronal sub-types will present different resiliences to different insults. Our studies were performed on glutamatergic hippocampal neurons, an extremely well-characterised model system for studying excitatory neurotransmission, and the most thoroughly neuronal type in terms of the physiology of small central synapses. Interestingly, transgenic mice overexpressing haSyn display the most robust overexpression in the hippocampus ^63^.

Future studies should further explore the hypothesis of specific functions of the protein by comparing the results to other types of synapses, in particular dopaminergic neurons, which are critically implicated in Parkinson’s disease. Although the synaptic transmission of dopaminergic neurons is less well characterised, new methods permitting the analysis of dopamine release at synapses, and a new focus on understanding the molecular physiology of dopaminergic synapses, are dramatically increasing our understanding of these structures ^64,65^.

aSyn is known to influence various cellular processes crucial for neuronal structure and function ^66^ and to interact with other proteins. Recent works have confirmed that aSyn interactions with both synapsin ^22^ and VAMP 2 ^59^ regulate aSyn function. These interactions with other molecules may induce synergistic effects or modulate aSyn’s effects and vary depending on the cellular context. In our experiments, the overexpression of haSyn did not lead to overall alterations in the synaptic marker proteins SNAP25 and PSD95. Interestingly, there was an increase in the size and intensity of PSD95-positive immunolabelled puncta, which may indicate a change in PSD-95 trafficking induced by haSyn overexpression in rat neurons. Importantly, however, our electrophysiological analyses in mouse neurons showed no evidence of altered postsynaptic function.

We also examined neuronal morphology in neurons overexpressing haSyn. Previous studies using animal models and cellular systems shown aSyn-associated alterations in neuronal morphology ^67–70^. In vitro studies using neuronal cultures overexpressing aSyn have revealed dendritic simplification, reduced dendritic spine density, and impaired neurite outgrowth compared to control neurons ^67^. Similarly, overexpression of aSyn in transgenic mice has been shown to induce neuritic pathology characterized by abnormal axonal swellings and dystrophic neurites ^68^ and compromised dendritic branching and less dendritic intersections ^70^. Notably, the dendritic arborization was preserved at an earlier time point of aSyn overexpression, indicating that aSyn produces progressive impairment of the dendritic arbor of neurons ^70^. However, other studies focused on dopaminergic neurons, using neuronal cultures and transgenic mice with aSyn overexpression, have shown that increased aSyn levels were not associated with significant structural abnormalities in dendritic or axonal compartments ^19,71^. Our experimental results did not reveal significant alterations in neuronal morphology following aSyn overexpression, which is moreover consistent with our observed lack of functional defects in neurons overexpressing haSyn.

These findings suggest that the effects of aSyn on neuronal structure may be nuanced and influenced by specific cellular contexts and may not necessarily result from elevated aSyn levels alone. While alterations in axonal or dendritic morphology have been observed in certain experimental paradigms involving aSyn overexpression, the general impact on neuronal morphology appears to be less pronounced compared to synaptic dysfunction. Therefore, further investigations into the intricate interactions involving aSyn and their consequences on cellular structures will enhance our understanding of synucleinopathies and may pave the way for innovative therapeutic strategies.

The diverse results observed in studies involving aSyn highlight the complex nature of this protein. Understanding the reasons behind these discrepancies is essential for developing effective therapeutic strategies for synucleinopathies. For that reason, it is crucial to consider factors such as expression levels, cellular context, experimental models, and protein interactions when interpreting and designing experimental approaches involving this protein. This highlighted the importance of having standard protocols in the field. Further investigations are needed to unravel the intricate mechanisms underlying aSyn-related neurodegeneration and to bridge the gap between in vitro findings and clinical relevance.

## ACKNOWLEDGEMENTS

This work was supported by the SFB1286 (B08 to. T.F.O. and A09 to Nils Brose). I.H.G-P. was supported by Fundación Mutua Madrileña. We thank the microscopy facility of the European Neuroscience Institute Göttingen (ENI) for their support with confocal microscopy and the Animal Facility of the Max Planck Institute for Multidisciplinary Sciences for the maintenance of mouse colonies. We thank Dr. Nils Brose, Max Planck Institute for Multidisciplinary Sciences, for providing project funding, suggestions, and support.

## AUTHOR CONTRIBUTIONS

J.D. and T.F.O. conceived and designed the experiments. P.I.S., performed experiments on rat primary cultures, I.H.G.-P., and J.D. performed experiments on mouse primary cultures. J.S.R., provided access to electrophysiology instrumentation, protocols and analysis tools. A.S., provided imaging support, and A.S., A.A.C., and S.O.R. provided analysis tools for immunohistochemistry data. P.I.S., I.H.G.-P., and J.D. analysed and interpreted the data. I.H.G.-P edited the data figures. P.I.S., I.H.G.- P., J.D. and T.F.O. wrote the manuscript. All authors revised and approved the manuscript.

## DATA AVAILABILITY

The datasets generated and/or analysed during the current study are available from the corresponding authors on request.

## FIGURE LEGENDS

**Supplementary Figure 1. Increased expression of aSyn at continental hippocampal mouse neurons infected with haSyn lentivirus.**

Primary hippocampal continental cultures from mice were infected with lentivirus mediating the expression of either eGFP alone or haSyn and eGFP. Representative immunoblots of human aSyn (A), total aSyn (B), and eGFP (C) (n=3) levels. (D) 1.82-fold increase in total aSyn levels in haSyn-infected neurons (n=3). β-tubulin was used as a loading control.

**Figure Supplementary 2. Lentivirus-mediated overexpression of haSyn increases PSD-95 fluorescence at postsynaptic sites**

(A-H) Representative images of immunocytochemistry of continental hippocampal neurons infected with eGFP or haSyn lentivirus against SNAP-25 (B,G) and MAP2 (C,H). Overlays are shown (A,F). Scale bars = 10 μm. Quantification of SNAP-25 synaptic mean grey intensity (D), synaptic area (E) and synaptic volume (I) (n=3). (J-K) Representative immunoblots (K) and respective quantification (J) of SNAP25 levels assessed in primary neurons transduced with lentivirus encoding for eGFP or haSyn for up to DIV21 (n=4-5). (L-U) Representative images of immunocytochemistry of continental hippocampal neurons infected with eGFP or haSyn lentivirus against PSD-95 (M,R) and MAP2 (N,S). Overlays are shown (L,Q). Scale bars = 10 μm. Quantification of PSD-95 synaptic mean grey intensity (O), synaptic area (P) and synaptic volume (T) (n=3). (U-V) Representative immunoblots (V) and respective quantification (U) of PSD95 levels assessed in primary neurons transduced with lentivirus encoding for eGFP or haSyn for up to DIV21 (n=4-5). Data are expressed as mean ± SEM; Student’s t-test; * p-value ≤ 0.05; *** p-value ≤ 0.001.

## REFERENCES

1. Maroteaux, L., Campanelli, J. T. & Scheller, R. H. Synuclein: A neuron-specific protein localized to the nucleus and presynaptic nerve terminal. J. Neurosci. (1988). doi:10.1523/jneurosci.08-08-02804.1988

2. Koss, D. J. et al. Nuclear alpha-synuclein is present in the human brain and is modified in dementia with Lewy bodies. Acta Neuropathol. Commun. (2022). doi:10.1186/s40478-022-01403-x

3. Maroteaux, L. & Scheller, R. H. The rat brain synucleins; family of proteins transiently associated with neuronal membrane. Mol. Brain Res. (1991). doi:10.1016/0169-328X(91)90043-W

4. Sharma, M. & Burré, J. α-Synuclein in synaptic function and dysfunction. Trends in Neurosciences (2023). doi:10.1016/j.tins.2022.11.007

5. Sulzer, D. & Edwards, R. H. The physiological role of α-synuclein and its relationship to Parkinson’s Disease. Journal of Neurochemistry (2019). doi:10.1111/jnc.14810

6. Perez, R. G. Editorial: The Protein Alpha-Synuclein: Its Normal Role (in Neurons) and Its Role in Disease. Frontiers in Neuroscience (2020). doi:10.3389/fnins.2020.00116

7. Spillantini, M. G., Crowther, R. A., Jakes, R., Hasegawa, M. & Goedert, M. alpha-Synuclein in filamentous inclusions of Lewy bodies from Parkinson’s disease and dementia with Lewy bodies (ubiquitinlsarkosyl-insoluble filamentslimmunoelectron microscopy). Neurobiol. Commun. by Max F. Perutz, Med. Res. Counc. (1998).

8. Spillantini, M. G. et al. α-synuclein in Lewy bodies. Nature 388, 839–40 (1997).

9. Outeiro, T. F. et al. Defining the Riddle in Order to Solve It: There Is More Than One “Parkinson’s Disease”. Movement Disorders (2023). doi:10.1002/mds.29419

10. Oliveira, L. M. A. et al. Alpha-synuclein research: defining strategic moves in the battle against Parkinson’s disease. npj Parkinson’s Disease (2021). doi:10.1038/s41531-021-00203-9

11. Chandra, S., et al. Double-knockout mice for alpha- and beta-synuclein: Effect on synaptic functions. PNAS (2004).

12. Greten-Harrison, B. et al. αβγ-Synuclein triple knockout mice revealage-dependent neuronal dysfunction. Proc. Natl. Acad. Sci. U. S. A. (2010). doi:10.1073/pnas.1005005107

13. Gureviciene, I., Gurevicius, K. & Tanila, H. Role of α-synuclein in synaptic glutamate release. Neurobiol. Dis. (2007). doi:10.1016/j.nbd.2007.06.016

14. Liu, S. et al. α-synuclein produces a long-lasting increase in neurotransmitter release. EMBO J. (2004). doi:10.1038/sj.emboj.7600451

15. Vargas, K. J. et al. Synucleins regulate the kinetics of synaptic vesicle endocytosis. J. Neurosci. (2014). doi:10.1523/JNEUROSCI.4787-13.2014

16. Vargas, K. J. et al. Synucleins Have Multiple Effects on Presynaptic Architecture. Cell Rep. (2017). doi:10.1016/j.celrep.2016.12.023

17. Singleton, A. B. et al. α-Synuclein Locus Triplication Causes Parkinson’s Disease. Science (80-. ). (2003). doi:10.1126/science.1090278

18. Nemani, V. M. et al. Increased Expression of α-Synuclein Reduces Neurotransmitter Release by Inhibiting Synaptic Vesicle Reclustering after Endocytosis. Neuron 65, 66–79 (2010).

19. Scott, D. A. et al. A pathologic cascade leading to synaptic dysfunction in α-synuclein-induced neurodegeneration. J. Neurosci. (2010). doi:10.1523/JNEUROSCI.1091-10.2010

20. Watson, J. B. et al. Alterations in corticostriatal synaptic plasticity in mice overexpressing human α-synuclein. Neuroscience (2009). doi:10.1016/j.neuroscience.2009.01.021

21. Logan, T., Bendor, J., Toupin, C., Thorn, K. & Edwards, R. H. α-Synuclein promotes dilation of the exocytotic fusion pore. Nat. Neurosci. (2017). doi:10.1038/nn.4529

22. Atias, M. et al. Synapsins regulate α-synuclein functions. Proc. Natl. Acad. Sci. U. S. A. (2019). doi:10.1073/pnas.1903054116

23. Villar-Piqué, A. et al. Environmental and genetic factors support the dissociation between α-synuclein aggregation and toxicity. Proc. Natl. Acad. Sci. U. S. A. (2016). doi:10.1073/pnas.1606791113

24. Tönges, L. et al. Alpha-synuclein mutations impair axonal regeneration in models of Parkinson’s disease. Front. Aging Neurosci. (2014). doi:10.3389/fnagi.2014.00239

25. Burgalossi, A. et al. Analysis of neurotransmitter release mechanisms by photolysis of caged Ca2+ in an autaptic neuron culture system. Nat. Protoc. (2012). doi:10.1038/nprot.2012.074

26. Jockusch, W. J. et al. CAPS-1 and CAPS-2 Are Essential Synaptic Vesicle Priming Proteins. Cell (2007). doi:10.1016/j.cell.2007.11.002

27. Paiva, I. et al. Alpha-synuclein deregulates the expression of COL4A2 and impairs ER-Golgi function. Neurobiol. Dis. (2018). doi:10.1016/j.nbd.2018.08.001

28. Barde, I., Salmon, P. & Trono, D. Production and Titration of Lentiviral Vectors. Curr. Protoc. Neurosci. (2010). doi:10.1002/0471142301.ns0421s53

29. Follenzi, A. & Naldini, L. Generation of HIV-1 derived lentiviral vectors. Methods Enzymol. (2002). doi:10.1016/S0076-6879(02)46071-5

30. Follenzi, A. & Naldini, L. HIV-based vectors. Preparation and use. Methods Mol. Med. (2002). doi:10.1385/1-59259-141-8:259

31. Ripamonti, S. et al. Transient oxytocin signaling primes the development and function of excitatory hippocampal neurons. Elife (2017). doi:10.7554/eLife.22466

32. Stevens, C. F. & Williams, J. H. Discharge of the readily releasable pool with action potentials at hippocampal synapses. J. Neurophysiol. (2007). doi:10.1152/jn.00857.2007

33. Rosenmund, C. & Stevens, C. F. Definition of the readily releasable pool of vesicles at hippocampal synapses. Neuron (1996). doi:10.1016/S0896-6273(00)80146-4

34. Longair, M. H., Baker, D. A. & Armstrong, J. D. Simple neurite tracer: Open source software for reconstruction, visualization and analysis of neuronal processes. Bioinformatics (2011). doi:10.1093/bioinformatics/btr390

35. Sato, Y. et al. Three-dimensional multi-scale line filter for segmentation and visualization of curvilinear structures in medical images. Med. Image Anal. (1998). doi:10.1016/S1361-8415(98)80009-1

36. Rhee, H. J. et al. An Autaptic Culture System for Standardized Analyses of iPSC-Derived Human Neurons. Cell Rep. (2019). doi:10.1016/j.celrep.2019.04.059

37. Daniel, J. A. et al. An intellectual-disability-associated mutation of the transcriptional regulator NACC1 impairs glutamatergic neurotransmission. Front. Mol. Neurosci. (2023). doi:10.3389/fnmol.2023.1115880

38. Crocker, J. C. & Grier, D. G. Methods of digital video microscopy for colloidal studies. J. Colloid Interface Sci. (1996). doi:10.1006/jcis.1996.0217

39. Rodríguez-Losada, N. et al. Overexpression of alpha-synuclein promotes both cell proliferation and cell toxicity in human SH-SY5Y neuroblastoma cells. J. Adv. Res. (2020). doi:10.1016/j.jare.2020.01.009

40. Winner, B. et al. In vivo demonstration that alpha-synuclein oligomers are toxic. PNAS 108, 4194–4199 (2011).

41. Ross, A. et al. Alleviating toxic α-Synuclein accumulation by membrane depolarization: Evidence from an in vitro model of Parkinson’s disease. Mol. Brain (2020). doi:10.1186/s13041-020-00648-8

42. Calabresi, P., Picconi, B., Parnetti, L. & Di Filippo, M. A convergent model for cognitive dysfunctions in Parkinson’s disease: the critical dopamine-acetylcholine synaptic balance. Lancet Neurology (2006). doi:10.1016/S1474-4422(06)70600-7

43. Kaufmann, W. E. & Moser, H. W. Dendritic anomalies in disorders associated with mental retardation. Cerebral Cortex (2000). doi:10.1093/cercor/10.10.981

44. Selkoe, D. J. Alzheimer’s disease is a synaptic failure. Science (2002). doi:10.1126/science.1074069

45. Stephan, K. E., Friston, K. J. & Frith, C. D. Dysconnection in Schizophrenia: From abnormal synaptic plasticity to failures of self-monitoring. Schizophrenia Bulletin (2009). doi:10.1093/schbul/sbn176

46. Yang, W. & Yu, S. Synucleinopathies: common features and hippocampal manifestations. Cellular and Molecular Life Sciences (2017). doi:10.1007/s00018-016-2411-y

47. Murthy, V. N., Schikorski, T., Stevens, C. F. & Zhu, Y. Inactivity produces increases in neurotransmitter release and synapse size. Neuron (2001). doi:10.1016/S0896-6273(01)00500-1

48. Murthy, V. N. & Stevens, C. F. Reversal of synaptic vesicle docking at central synapses. Nat. Neurosci. (1999). doi:10.1038/9149

49. Imig, C. et al. The Morphological and Molecular Nature of Synaptic Vesicle Priming at Presynaptic Active Zones. Neuron (2014). doi:10.1016/j.neuron.2014.10.009

50. Chopra, A., Lang, A. E., Höglinger, G. & Outeiro, T. F. Towards a biological diagnosis of PD. Parkinsonism and Related Disorders (2024). doi:10.1016/j.parkreldis.2024.106078

51. Brás, I. C. et al. Synucleinopathies: Where we are and where we need to go. Journal of Neurochemistry (2020). doi:10.1111/jnc.14965

52. Outeiro, T. F. et al. Dementia with Lewy bodies: An update and outlook. Molecular Neurodegeneration (2019). doi:10.1186/s13024-019-0306-8

53. Anwar, S. et al. Functional alterations to the nigrostriatal system in mice lacking all three members of the synuclein family. J. Neurosci. (2011). doi:10.1523/JNEUROSCI.6194-10.2011

54. Abeliovich, A. et al. Mice lacking α-synuclein display functional deficits in the nigrostriatal dopamine system. Neuron 25, 239–252 (2000).

55. Burré, J. et al. α-Synuclein promotes SNARE-complex assembly in vivo and in vitro. Science (80-. ). 329, 1663–1667 (2010).

56. Diao, J. et al. Native α-synuclein induces clustering of synaptic-vesicle mimics via binding to phospholipids and synaptobrevin-2/VAMP2. Elife 1–17 (2013). doi:10.7554/eLife.00592

57. Janezic, S. et al. Deficits in dopaminergic transmission precede neuron loss and dysfunction in a new Parkinson model. Proc. Natl. Acad. Sci. U. S. A. (2013). doi:10.1073/pnas.1309143110

58. Dewitt, D. C. & Rhoades, E. α-synuclein can inhibit SNARE-mediated vesicle fusion through direct interactions with lipid bilayers. Biochemistry (2013). doi:10.1021/bi4002369

59. Sun, J. et al. Functional cooperation of α-synuclein and VAMP2 in synaptic vesicle recycling. Proc. Natl. Acad. Sci. U. S. A. (2019). doi:10.1073/pnas.1903049116

60. Junge, H. J. et al. Calmodulin and Munc13 form a Ca2+ sensor/effector complex that controls short-term synaptic plasticity. Cell (2004). doi:10.1016/j.cell.2004.06.029

61. López-Murcia, F. J., Reim, K., Jahn, O., Taschenberger, H. & Brose, N. Acute Complexin Knockout Abates Spontaneous and Evoked Transmitter Release. Cell Rep. (2019). doi:10.1016/j.celrep.2019.02.030

62. Steidl, J. V., Gomez-Isla, T., Mariash, A., Ashe, K. H. & Boland, L. M. Altered short-term hippocampal synaptic plasticity in mutant α-synuclein transgenic mice. Neuroreport (2003). doi:10.1097/00001756-200302100-00012

63. Yavich, L., Jäkälä, P. & Tanila, H. Abnormal compartmentalization of norepinephrine in mouse dentate gyrus in α-synuclein knockout and A30P transgenic mice. J. Neurochem. (2006). doi:10.1111/j.1471-4159.2006.04098.x

64. Cai, X. et al. Dopamine dynamics are dispensable for movement but promote reward responseso Title. Nature 635, 406–414 (2024).

65. Elizarova, S. et al. A fluorescent nanosensor paint detects dopamine release at axonal varicosities with high spatiotemporal resolution. Proc. Natl. Acad. Sci. U. S. A. (2022). doi:10.1073/pnas.2202842119

66. Lee, H. J., Patel, S. & Lee, S. J. Intravesicular localization and exocytosis of α-synuclein and its aggregates. J. Neurosci. (2005). doi:10.1523/JNEUROSCI.0692-05.2005

67. Dauer, W. & Przedborski, S. Parkinson’s disease: Mechanisms and models. Neuron (2003). doi:10.1016/S0896-6273(03)00568-3

68. Lee, V. M. Y. & Trojanowski, J. Q. Mechanisms of Parkinson’s Disease Linked to Pathological α-Synuclein: New Targets for Drug Discovery. Neuron (2006). doi:10.1016/j.neuron.2006.09.026

69. Eslamboli, A. et al. Long-term consequences of human alpha-synuclein overexpression in the primate ventral midbrain. Brain (2007). doi:10.1093/brain/awl382

70. Ledonne, A. et al. Morpho-Functional Changes of Nigral Dopamine Neurons in an α-Synuclein Model of Parkinson’s Disease. Mov. Disord. (2023). doi:10.1002/mds.29269

71. Yavich, L., Tanila, H., Vepsäläinen, S. & Jäkälä, P. Role of α-synuclein in presynaptic dopamine recruitment. J. Neurosci. (2004). doi:10.1523/JNEUROSCI.2559-04.2004

