## Supplementary figures and images for "Glutamatergic synaptic resilience to overexpressed human alpha-synuclein"

### Supplementary Figure 1

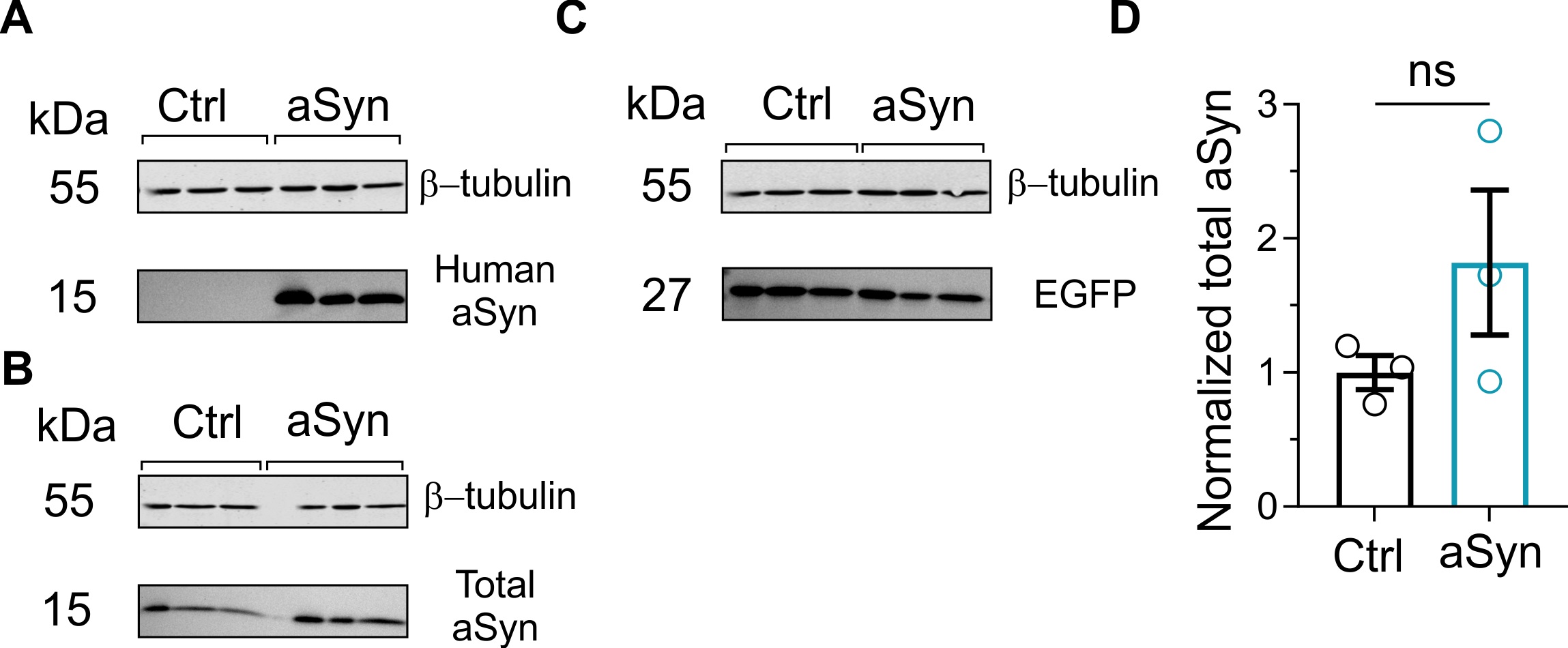

### Supplementary Figure 2

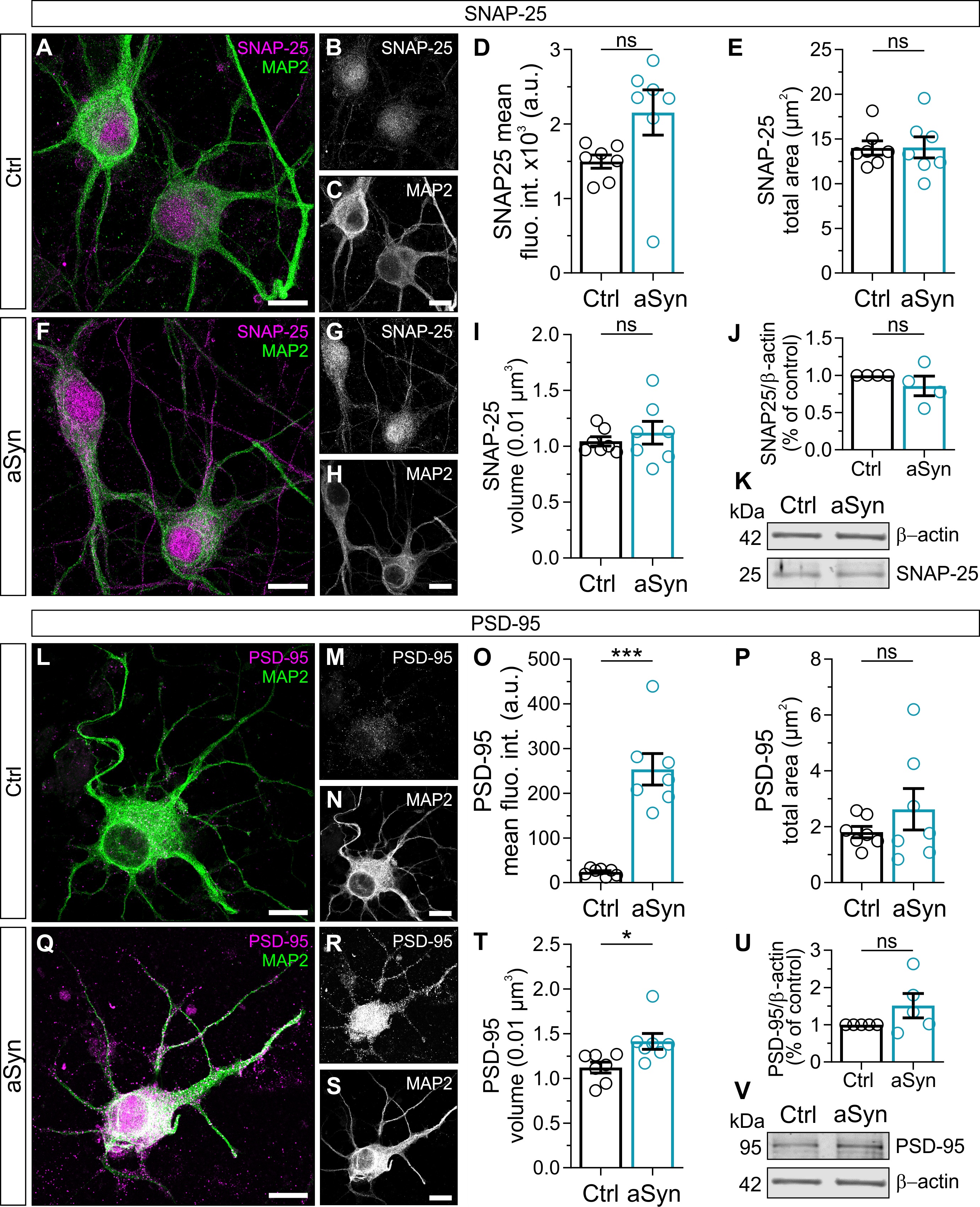
